# *De novo* mutations implicate novel genes with burden of rare variants in Systemic Lupus Erythematosus

**DOI:** 10.1101/139238

**Authors:** Venu Pullabhatla, Amy L. Roberts, Myles J. Lewis, Daniele Mauro, David L. Morris, Christopher A. Odhams, Philip Tombleson, Ulrika Liljedahl, Simon Vyse, Michael A. Simpson, Sascha Sauer, Emanuele de Rinaldis, Ann-Christine Syvänen, Timothy J. Vyse

## Abstract

The omnigenic model of complex diseases stipulates that the majority of the heritability will be explained by the effects of common variation on genes in the periphery of core disease pathways. Rare variant associations, expected to explain far less of the heritability, may be enriched in core disease genes and thus will be instrumental in the understanding of complex disease pathogenesis and their potential therapeutic targets. Here, using complementary whole-exome sequencing (WES), high-density imputation, and *in vitro* cellular assays, we identify three candidate core genes in the pathogenesis of Systemic Lupus Erythematosus (SLE). Using extreme-phenotype sampling, we sequenced the exomes of 30 SLE parent-affected-offspring trios and identified 14 genes with missense *de novo* mutations (DNM), none of which are within the >80 SLE susceptibility loci implicated through genome-wide association studies (GWAS). In a follow-up cohort of 10,995 individuals of matched European ancestry, we imputed genotype data to the density of the combined UK10K-1000 genomes Phase III reference panel across the 14 candidate genes. We identify a burden of rare variants across *PRKCD* associated with SLE risk (*P*=0.0028), and across *DNMT3A* associated with two severe disease prognosis sub-phenotypes (*P*=0.0005 and *P*=0.0033). Both genes are functional candidates and significantly constrained against missense mutations in gene-level analyses, along with *C1QTNF4*. We further characterise the TNF-dependent functions of candidate gene *C1QTNF4* on NF-κB activation and apoptosis, which are inhibited by the p.His198Gln DNM. Our results support extreme-phenotype sampling and DNM gene discovery to aid the search for core disease genes implicated through rare variation.

**Significance Statement:** Rare variants, present in <1% in population, are expected to explain little of the heritability of complex diseases, such as Systemic Lupus Erythematosus (SLE), yet are likely to identify core genes crucial to disease mechanisms. Their rarity, however, limits the power to show their statistical association with disease. Through sequencing the exomes of SLE patients and their parents, we identified non-inherited *de novo* mutations in 14 genes and hypothesised that these are prime candidates for harbouring additional disease-associated rare variants. We demonstrate that two of these genes also carry a significant excess of rare variants in an independent, large cohort of SLE patients. Our findings will influence future study designs in the search for the ‘missing heritability’ of complex diseases.

## Introduction

Considerable progress has been made in elucidating the genetic basis of complex diseases. The vast majority of identified disease-associated genetic polymorphisms are common in the population and the risk alleles impart a modest individual increment to the likelihood of developing disease. Although large-scale genome-wide association studies (GWAS) have so far explained less of the heritability than originally predicted (1), much of the ‘missing heritability’ is expected to be accounted for by common variants with effect sizes below the genome-wide significance threshold (2). However, under the newly proposed omnigenic model of complex traits, the majority of associated common variants – both identified and unidentified - will primarily be found in periphery genes expressed in relevant cell types but not necessarily biologically relevant to disease (3).

In contrast, the role of rare variants in complex disease is largely unknown and often dismissed. A recent study, however, with an extremely large sample size, identified rare and low frequency variants contributing to the genetic variance of adult human height (4) – a polygenic trait with a genetic architecture similar to that of complex diseases (5) - suggesting previous complex disease studies with seemingly large sample sizes were perhaps still insufficiently powered to detect rare variant associations (6). Furthermore, studies of rare variants typically find gene sets enriched in biologically relevant functions/pathways (3, 7, 8). Therefore, although estimated to explain less of the heritable disease risk at a population level than common variants, identifying rare and low frequency variants is of paramount importance to understanding disease pathogenesis as they are likely to implicate biologically relevant core genes (3). Supporting the theory that common and rare variant associations will be found in discrete gene sets is the lack of additional rare variant associations in GWAS genes (9).

Exome-wide searches, which provides a highly enriched source of potential disease-causing mutations (10), have revealed limited numbers of rare variation associated with complex diseases. Even though greater statistical power is achieved by gene-level analyses whereby aggregated variants are tested for an allelic burden of collective rare variation, widely used gene-based association tests have been shown to lack power at the exome-wide level (11). Coupled with the insufficient sample sizes currently available in the study of most complex diseases, hypothesis-free searches for core genes with rare variant associations are unlikely to be fruitful.

Our strategy to address this problem in autoimmune disease Systemic Lupus Erythematosus (SLE), is outlined here and summarised in Fig. 1. Using a discovery cohort of 30 unrelated SLE cases with a severe disease (young age of onset and clinical features associated with poorer outcome), we hypothesized that these individuals would exhibit unique mutation events in their protein-coding DNA that may predisposed to disease risk. We undertook whole exome sequencing (WES) in 30 family trios (both parents and affected offspring) and scrutinized the data for non-inherited *de novo* mutations (DNM) in the individual with SLE to identify a group of candidate genes for an independent follow-up rare variant analysis. This method allowed the identification of novel loci harbouring disease risk through collective rare variation, and emphasises the value of phenotypic extremes in the search for core genes in multifactorial disorders (12).

**Figure 1.**
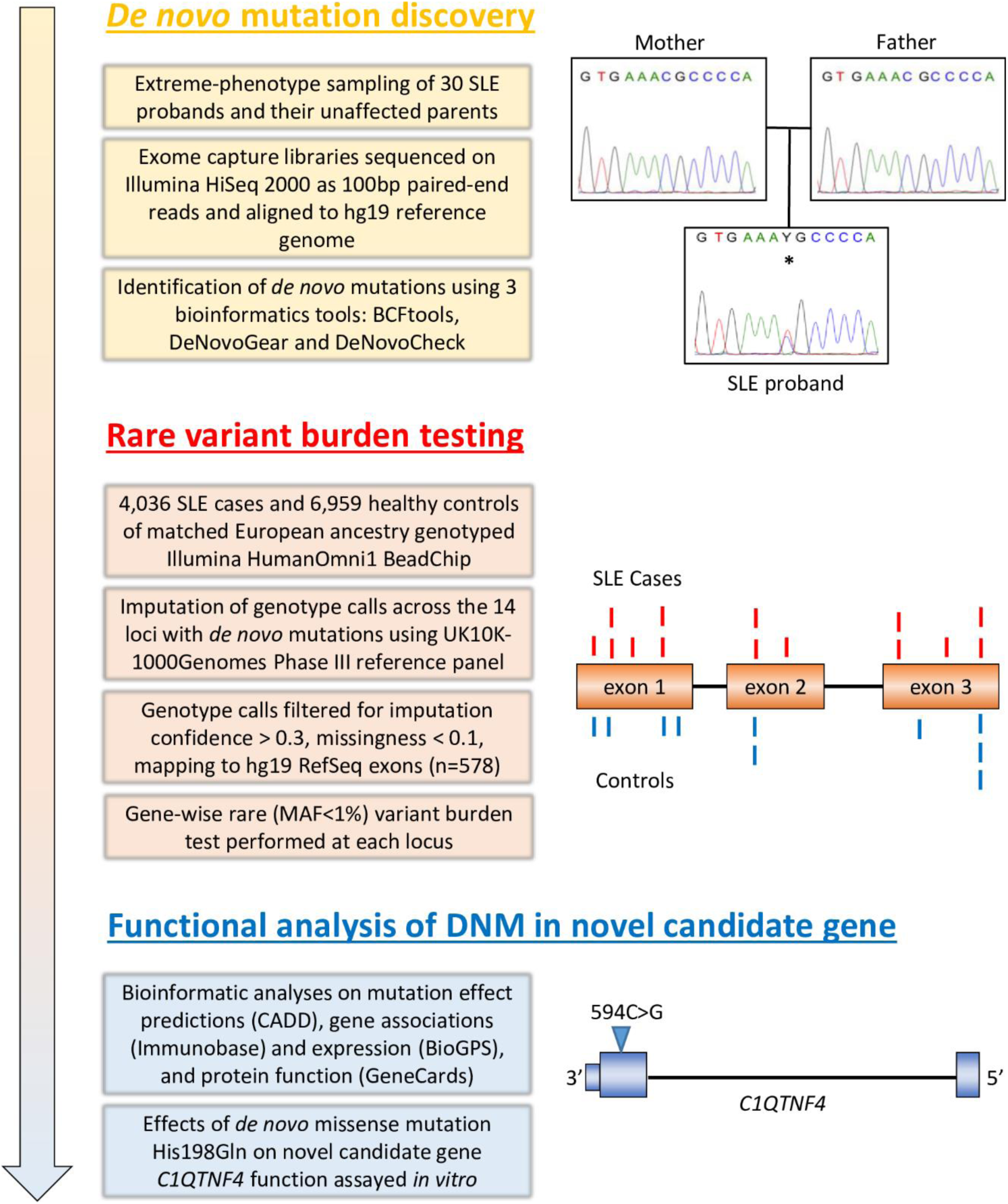
Overview of study. *De novo* mutations (DNM) in a discovery cohort revealed candidate genes for imputation-based rare variant burden testing using a follow-up cohort. Independent functional analyses demonstrate the functional effects of one DNM in a candidate gene.

## Results

### Identification of DNM in extreme-phenotype SLE cases

We screened for DNM by WES of 30 family trios with an affected offspring with more severe SLE (Fig. S1). A total of 584,798 variants (≥20X), including single nucleotide variants and indels, were identified in the 30 affected probands. Using three bioinformatic tools and employing conservative parameters, 17 putative missense DNM were identified across 17 genes (Table S1; Fig. S2). We also analysed the SLE proband WES data alone, without the unaffected parents. This revealed 1,194 non-silent, heterozygous, rare variants in 1,067 genes distributed across the genome, which would make prioritisation for downstream analysis a difficult task, highlighting the benefit of parent-offspring trio sequencing (Fig. S3). Sanger sequencing confirmed 14 true positive non-silent DNM (Table 1; Table S2), present in the SLE proband but absent in both parents and any unaffected siblings, in 11 of the 30 probands (36.7%) for further analysis. No DNM was found in any of the >80 known SLE-associated genes. Of the three false positive DNM (11.7%; Table S1) one, within *LAMC2*, is likely a result of germline mosaicism because, although not observed in either parent, it is observed in an unaffected sibling in addition to the SLE proband (13), and the other two variants are within *KRTAP10-2* and *KLRC1* - both members of highly homologous gene families. Such sequence identity may have caused false positive identification of DNM in the WES analysis and suggests our NGS error-prone genes (NEPG) filter, which removes loci known to be problematic for genome mapping during NGS analyses, should have been more conservative. Indeed the KRLC1 p.Ile225Met missense variant appears to be a polymorphic Paralogous Sequence Variant (PSV) – the paralogous variant being Met223Ile in KLRC2.

**Table 1:**
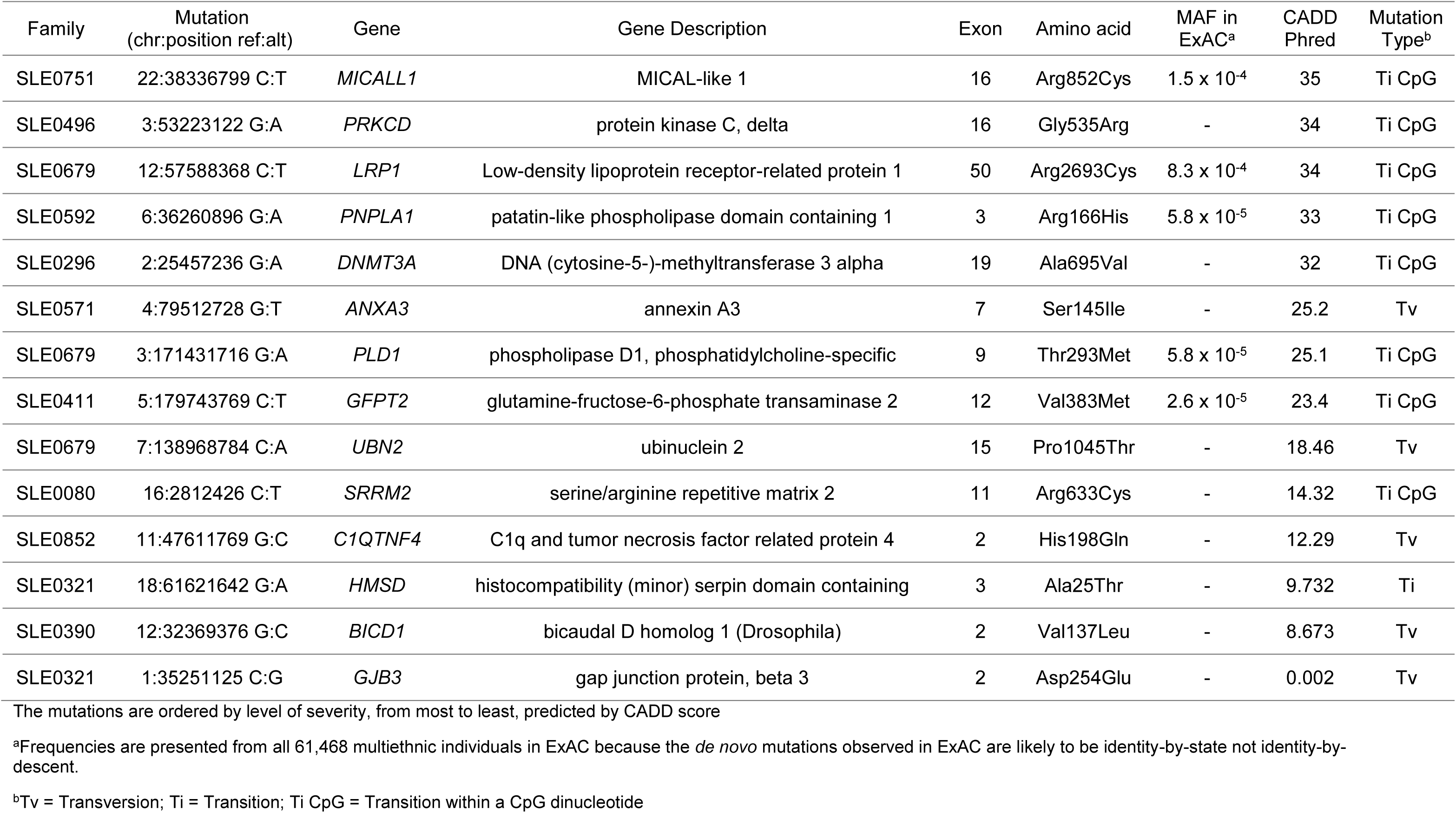
*De novo* mutations in SLE probands with extreme phenotypes

### Variant- and gene-level functional characterisation of DNM

In order to best predict the phenotypic effect of the 14 DNM, we used both variant-level and gene-level metrics (14). We used the ExAC database (15) and Combined Annotation Dependent Depletion (CADD) scores (16) to characterise the frequency and predicted functional effects, respectively, of the variants. Five of the 14 DNM – found in *MICALL1*, *LRP1*, *PNPLA1*, *PLD1*, and *GFTP2* - have been observed, at very rare frequencies, in the ∼60,000 exomes documented in ExAC (Table 1). All five mutations are CpG transitions and therefore likely to be identity-by-state, reflecting the higher mutability rate of these sites. Within the mutation set, five (35.7%) – found in *DNMT3A, PRKCD, MICALL1, LRP1,* and *PNPLA1* – have CADD Phred scores >30, placing them in the top 0.1% of possible damaging mutations in the human genome (Table 1). We further explored the function, expression (BioGPS), existing autoimmunity associations (ImmunoBase), and gene-level constraint against missense mutations (ExAC), of the DNM genes to build a profile of *a priori* evidence of a role in SLE pathogenesis. None of the candidate genes have been previously associated with SLE through GWAS in any population (17). We also identify candidate genes through known/predicted function and expression profiles (*C1QTNF4, SRRM2, HMSD*), and four genes (*PRKCD, DNMT3A, C1QTNF4* and *LRP1)* with a significant (Z>3.09) constraint against missense variants (Table 2). However, across the entire gene set, there was no difference in the median Z-score (0.50) compared with the median Z-score across all genes in ExAC (0.51).

**Table 2.** Evidence for role of *de novo* mutation gene in autoimmunity

| Gene | Functional Candidate<sup>a</sup> | Association with SLE<sup>b</sup> | Associations with other AID<sup>b</sup> | Immune cell type with highest expression<sup>c</sup> | Missense Constraint<sup>d</sup> |
| --- | --- | --- | --- | --- | --- |
| *PRKCD* | B cell signaling and self-antigen induced B cell tolerance induction | Monogenic forms<sup>30</sup> | IBD, UC, CD<sup>28</sup> | Dendritic | 3.75* |
| *DNMT3A* | DNA methyltransferase | Candidate gene study<sup>35</sup> | CD<sup>29</sup> | - | 4.31* |
| *C1QTNF4* | Pro-inflammatory cytokine | - | - | CD34+ | 3.17* |
| *SRRM2* | Spliceosome-associated pre-mRNA splicing | - | - | CD8+ | No data |
| *LRP1* | Endo/Phagocytosis of apoptotic cells | - | - | - | 10.60* |
| *HMSD* | Minor histocompatibility antigen | - | - | n/a | 0.25 |
| *UBN2* | DNA binding | - | - | - | 0.01 |
| *ANXA3* | - | - | RA<sup>17</sup> | - | -0.37 |
| *PLD1* | - | - | - | Lymphoblasts | -0.73 |
| *PNPLA1* | - | - | - | - | 0.27 |
| *GFPT2* | - | - | - | - | 1.59 |
| *BICD1* | - | - | - | - | 2.12 |
| *GJB3* | - | - | - | - | -0.81 |
| *MICALL1* | - | - | - | - | 0.50 |

Genes appear in descending order of supporting evidence. UC=ulcerative colitis, CD=Crohn's Disease, IBD=inflammatory bowel disease, RA=Rheumatoid Arthritis
<sup>a</sup> See Table S5
<sup>b</sup> See Table S6
<sup>c</sup> See SFigure S4. Data from BioGPS. If gene expression is highest in immune cells compared to all other cells, the immune cell type with highest expression is listed.
<sup>d</sup> Gene-wise ExAC Constraint Z-scores. Genes with significant restraint against missense variants are highlighted with an asterisk.

### *PRKCD* and *DNMT3A* as novel SLE genes

Although the variant- and gene-level metric analyses suggested intriguing functional candidates, we took a comprehensive approach and tested each locus for an allelic burden of rare variation. We hypothesised that, while some observed DNM were random background variation as present in the exome of every individual regardless of disease status (18), others may be reflecting a hitherto unknown gene contributing to SLE risk, and this may be shown through rare variant burden.

Therefore, genotype data was imputed (Fig. S6 and S7) to the density of the combined UK10K and 1000 genomes Phase III reference panel (UK10K-1000GP3) across all 14 DNM genes in a follow-up cohort of 10,995 individuals of matched European ancestry previously genotyped on the Illumina HumanOmni1 BeadChip (19). Under the hypothesis that rare variants at these loci would be causal and not protective, we employed a one-tailed collapsing burden test (20) to survey each of the 14 genes for an excess of aggregated rare (MAF<1%) exonic variants in SLE cases compared with healthy controls. We identify an association of *PRKCD* rare variants with SLE (Table S3; *P*=0.0028; n_cases_=4,036). In sub-phenotype analyses, we identify collective rare exonic variants in *DNMT3A* associated with both anti-dsDNA (Table S3; *P*=0.0005; n_cases_=1,261) and renal involvement with hypocomplementemia (Table S3; *P*=0.0033; n_cases_=186), both of which are markers of more severe disease. We also collapsed all exons from the 14 genes together to test for an overall burden of rare variants across these loci. These analyses revealed no excess of rare exonic variants across the grouped genes, reflecting the hypothesis that some/most genes will not be relevant to disease status because the observed DNM are random background variation only. These data reflect the results of our gene-level constraint metric, in which the aggregated gene set do not have a significant mutation constraint. Together these results suggest further prioritisation based on gene-level metrics would not have resulted in true positive associations being excluded from analyses.

### Effect of DNM p.His198Gln on C1QTNF4 function

Although no rare variant association was found at the novel candidate gene *C1QTNF4*, it’s potential role in disease is supported by gene-level metrics – it is a compelling functional candidate and one of four genes constrained against missense variants (ExAC gene-level constraints Z=3.17, Table 2). Although gene coding length does not correlate with missense constraint scores (15), the small (<1Kb) coding sequence of this candidate gene may have contributed to insufficient power to detect a rare variant association in the burden testing. On the variant-level, the DNM in *C1QTNF4* generates a p.His198Gln sequence change with a modest CADD score of 12.3 (Table 1). Although useful in the absence of suitable functional assays, the sensitivity of bioinformatic prediction tools is known to be suboptimal. Where functional assays are available, previous studies have also demonstrated functional effects of variants predicted to be tolerated/benign (21). We therefore pursued a functional analysis of the p.His198Gln DNM detected in the *C1QTNF4* gene as an alternative method to add support for its potential role in disease. Although its function is rather poorly understood, the protein product, C1QTNF4 (CTRP4) is secreted and may act as a cytokine, as it has homology with TNF and the complement component C1q (Fig. 2). C1QTNF4 has been shown to influence NF-κB activation (22), a pathway known to be implicated in SLE pathogenesis, therefore we looked for an effect of the p.His198Gln mutation on NF-κB production. Using a HEK293-NF-κB reporter cell line, we showed that C1QTNF4 p.His198Gln mutant protein was expressed and that it inhibited the NF-κB activation generated by exposure to TNF (Fig. 2). Furthermore, we showed that the fibroblast L929 cell line, which is sensitive to TNF-induced cell death, was rescued by exposure to C1QTNF4 p.His198Gln, but not by wild type C1QTNF4. Thus, the mutant form of C1QTNF4 appears to inhibit some of the actions of TNF (23–25).

**Figure 2.**
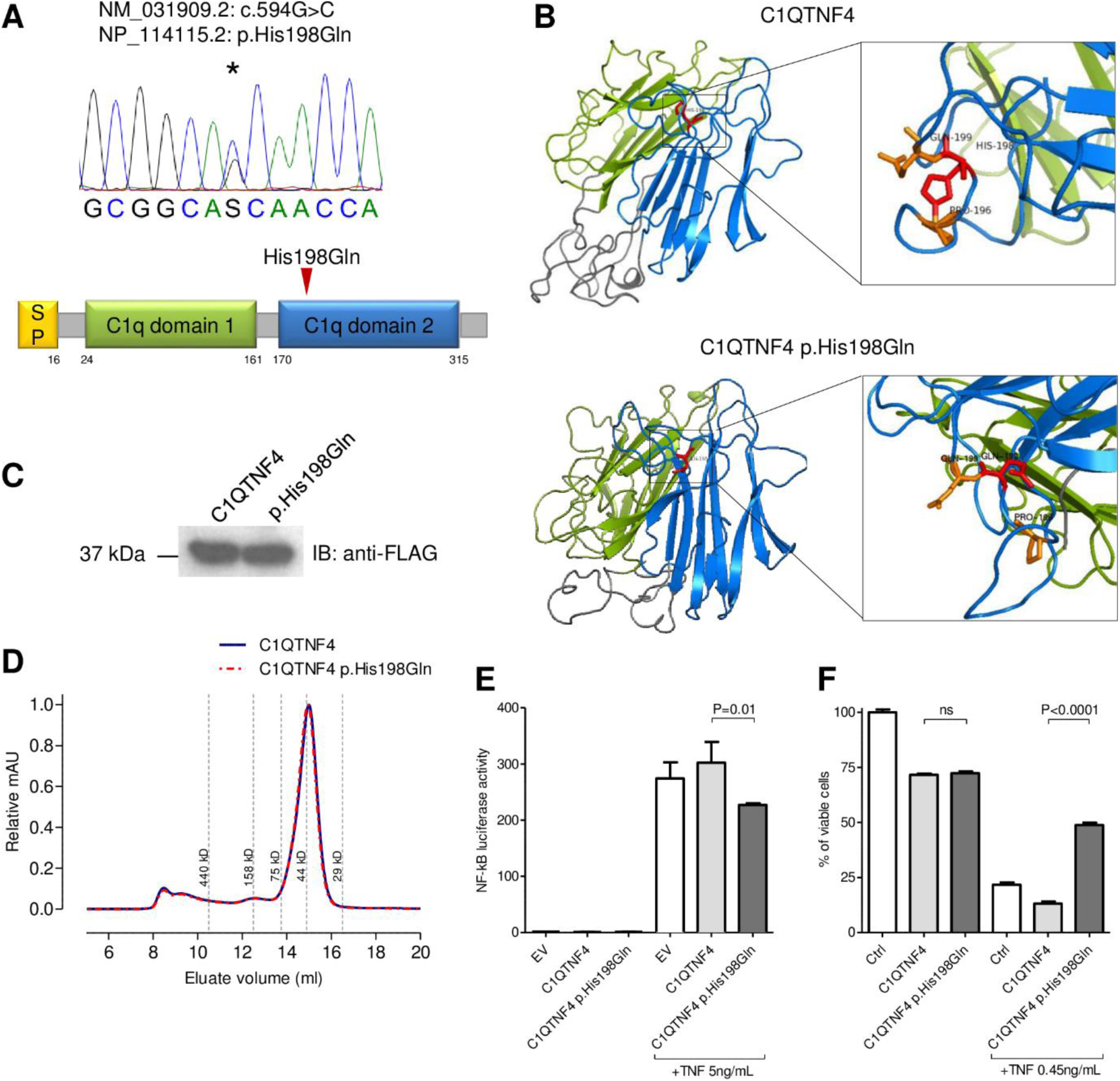
Structural and functional characterization of C1QTNF4 p.His198Gln substitution. (A) Domain organization of human C1QTNF4, showing signal peptide (yellow), first C1q domain (green), second C1q domain (blue) and linker peptides (grey). Arrow highlights substitution site. (B) 3D structure prediction of C1QTNF4 and C1QTNF4 p.His198Gln using Phyre2 (47). Ribbons show the interaction between the positively charged Histidine 198 and Proline 196 lost in C1QTNF4 p.His198Gln due to the substitution of Histidine with Glutamine. (C) Immunoblot demonstrating that p.His198Gln does not affect secretion of C1QTNF4 in HEK293 supernatants. (D) Size exclusion chromatography profile showing no difference in oligomerisation between supernatant containing C1QTNF4 (blue) and C1QTNF4 p.His198Gln (red). (E) Luciferase assay in HEK293-NF-κB reporter cell line showing that C1QTNF4 p.His198Gln inhibits NF-κB activation in response to 4h stimulation with 5ng/mL TNFα. Error bars represent standard error of the mean. (F) Inhibition of L929 induced cell death by C1QTNF4 p.His198Gln after 24h of stimulation with 0.45 ng/mL TNFα in presence of Actinomycin 1μg/ml. EV=empty vector.

### DNM genes do not harbour common variant associations

We next tested for additional common variant associations at these 14 loci using the high-density UK10K-1000GP3 imputed data. No significant association at any locus was observed with overall risk in a case-control comparison, nor with anti-dsDNA (n_cases_=1,261) or renal-involvement with hypocomplementemia (n_cases_=186) sub-phenotypes (Table S4). The lack of an associated common variant within *PRKCD* and *DNMT3A* supports the hypothesis that discrete gene sets will be identified through rare and common variant associations, with the former expecting to be enriched for core disease genes (3).

## Discussion

To fully understand the pathogenesis of complex diseases we must analyse the full frequency spectrum of genetic variants (4). The study of rare variants associated with disease is of paramount importance to the discovery of core genes that have the potential to be therapeutic targets (12). Our data support the omnigenic hypothesis that rare genetic risk may be found in a discrete set of non-canonical susceptibility genes, as we report an association of collective rare variation across *PRKCD* and *DNMT3A*, and found no evidence of an association with common variants across these loci. This, to the best of our knowledge, is the first WES study in polygenic cases of autoimmune disease to use DNM discovery to identify candidate genes for rare variant analyses. Furthermore, our study supports the importance of phenotypic extremes in elucidating the genetic basis of multifactorial disorders (26).

Searching GWAS-identified canonical disease susceptibility genes for additional rare variant risk has not been fruitful. Although there are examples – and perhaps more to discover – of canonical disease genes harbouring both common and rare risk alleles (27), the vast majority of such loci do not. Indeed the common variant associated loci which have also been shown to harbor rare coding variant risk are often those distinct minority of loci where the common polymorphisms are non-silent coding variants (e.g. *NCF2* (9)). It is important to note, however, that the separation of periphery and core genes may not necessarily be binary (3).

*DNMT3A* and *PRKCD*, although hitherto not associated with polygenic SLE, are known autoimmunity susceptibility loci; *DNMT3A* is associated with Crohn’s disease (CD) (28) and *PRKCD* is associated with both CD and ulcerative colitis (UC) (29). The notion that a locus could harbour common variants contributing to one autoimmune disease and rare variants contributing to another is intriguing, and could provide further hypothesis-driven searches in the hunt for disease-specific core genes.

A functional missense variant p.G510S (c.G1528A) in *PRKCD* has previously been reported in a consanguineous family with monogenic SLE (30). It was demonstrated that the *PRKCD*-encoded protein, PRCδ, was essential in the regulation of B cell tolerance and affected family members with the homozygous mutation had increased numbers of immature B cells. Our study implicates the role of rare variants in *PRKCD* in the broader context of SLE susceptibility, beyond a monogenic recessive disease model. Indeed the analysis of rare and low frequency variants contributing to human height found significant overlap with genes mutated in monogenic growth disorders (4). Furthermore, *PRKCB*, another member of the protein kinase C gene family, has been implicated in SLE risk in a Chinese study (31).

*DNMT3A*, a DNA methyltransferase, is a very intriguing candidate gene for SLE as altered patterns of DNA methylation are reported in autoimmune diseases (32), and hypomethylation of apoptotic DNA has been reported to induce autoantibody production in SLE (33). DNA methylation changes are also associated with monozygotic twin discordance in SLE (34). A candidate gene study previously reported a trend of association between the common *DNMT3A* intronic SNP rs1550117 (MAF∼7%) and SLE in a European cohort (35). Our analysis did not replicate this finding (*P*=0.23) and found no evidence of a common variant association at this locus. Instead we find an association of collective rare variants and SLE sub-phenotypes and emphasises the importance of deep phenotyping and the potential role of rare variants in specific sub-phenotype, or indeed autoimmune, manifestations. Despite progress with diagnosis and treatment, particular SLE sub-phenotypes – including those used in this study - are still associated with reduced life expectancy. Therefore, elucidating the specific underlying genetic risk is of paramount importance.

Through two in vitro assays, we demonstrated the functional effect of a DNM, p.His198Gln in *C1QTNF4*, despite this mutation being predicted to be of little functional importance across variant-level prediction tools. We showed the mutated protein product of *C1QTNF4*, C1QTNF4, inhibits some TNF-mediated cellular responses, including activation of NF-κB and TNF-induced apoptosis. The role of TNF in SLE is complex and incompletely understood, although, in this context, it is noteworthy that TNF inhibition may promote antinuclear autoimmunity (24). Gene-level metrics for *C1QTNF4* were supportive of a role in disease and our result support the importance of combined gene- and variant-level metrics, and the dangers of relying heavily on variant-level metrics alone, when interpreting the potential role of mutations (14). *C1QTNF6* is a known susceptibility locus for Type 1 Diabetes and is implicated in Rheumatoid Arthritis (36, 37), and an association with SLE has recently been in a transancestral Immunochip analysis (38). Together these data suggest a potential role of the hitherto understudied *C1QTNF* superfamily of genes in autoimmunity.

Although our study allowed a comprehensive approach to test all DNM genes for allelic burden of rare variants, our results show that filtering based on gene- or variant-level metrics would not have resulted in true associations of *DNMT3A* and *PRKCD* being missed. When larger datasets require further prioritisation of genes, we suggest both variant- and gene-level metrics are used.

Each human - regardless of the disease status - is estimated to have one DNM in their exome (18). The simple presence of a provisionally functional DNM in a proband is therefore not sufficient evidence that it contributes to disease risk. A major challenge of WES studies, therefore, is how to differentiate between variants truly important to disease and background variation (39). In light of recent studies which have demonstrated the limitations of large-scale exome-wide case-control studies in detecting rare variant associations (6, 40), despite such associations being found when no limitation on sample size exists (4), our results support extreme-phenotype sampling and DNM discovery to aid a hypothesis-driven search for rare variant associations with complex diseases, in the hunt to determine core disease genes.

## Methods

### Selection of trios for sequencing

SLE patients of European ancestry – as determined by genome-wide genotyping as part of a GWAS (19) - were selected from the UK SLE genetic repository assembled in the Vyse laboratory on the following criteria: age of onset of SLE < 25 years (median age 21 years); more marked disease phenotype as shown by either evidence for renal involvement as per standard classification criteria and/or the presence of hypocomplementemia and anti-dsDNA autoantibodies; and DNA available from both unaffected parents. The 30 trios (90 individuals) were exome sequenced, as described in SI Methods. Ethical approval for the research was granted by the NRES Committee London (12/LO/1273 and 06/MRE02/9).

### DNM calling

Three bioinformatics tools with conservative parameters were used for DNM screening: BCFtools (41), DeNovoGear (42) and DeNovoCheck (43). A detailed description of the methods applied can be found in SI Methods. Briefly, 454 variants were identified with BCFtools and DeNovoGear and eight additional variants were identified by DeNovoCheck and validated by IGV, resulting in a total of 462 variants, which map to 257 genes. The variants were next filtered sequentially filtered (Fig. S2): (A) Removal of NGS error prone genes (NEPG); (B) Fulfil a Het:Ref:Ref for Child:Father:Mother *de novo* pattern of inheritance and further selected variants that did not contain any trace of alternate allele in any of the parents; (C) Non-silent variant annotation. This process resulted in a total of 17 variants in 17 genes (Table S1).

### Analysis of whole exome sequencing (WES) in cases only

584,798 variants with ≥20X coverage depth and within Gencode capture regions were identified in the analysis of 30 SLE probands only. Stringent filters were applied for variant refinement, described in full in SI Methods, resulting in 1194 variants in 1067 genes (Fig.S3).

### Sanger Sequencing confirmation

Primers were designed using Primer 3. 10ng of DNA from SLE probands, any unaffected siblings and both parents was amplified with Hot Start Taq polymerase. PCR products were first purified with EXO-SAP before BigDye labelling in a linear PCR and sequenced on an ABI 3300XL. Primers and PCR conditions available on request. The reads were analysed using Chromas Lite (v.2.1.1)

### Imputation

Illumina HumanOmni1 BeadChip genotype data from 6,995 controls and 4,036 SLE patients of matched European ancestry were used, which had undergone quality control as previously described including Principal Component Analysis (PCA) to account for population structure (19).The UK10K (REL-2012-06-02) plus 1000 Genomes Project Phase3 data (release 20131101.v5) merged reference panel (UK10K-1000GP3) was accessed through the European Genome-phenome Archive (EGAD00001000776). The genotype data were imputed using the UK10K-1000GP3 reference panel across the coding regions of the 14 DNM genes plus a 2Mb flanking region. To increase the accuracy of imputed genotype calls, a full imputation without pre-phasing was conducted using IMPUTE2 (44, 45). Imputed genotypes were filtered for confidence using an info score (IMPUTE2) threshold of 0.3 (Fig. S6 and S7). The most likely genotype from IMPUTE2 was taken if its probability was > 0.5. If the probability fell below this threshold, it was set as missing. Variants with >10% missing genotype calls were removed for further analysis. All individuals had <8% missing genotype data.

### Rare variant burden tests

Imputed data were filtered, using Plink v1.9, to include only variants mapping to coding exons of hg19 RefSeq transcripts. Plink/SEQv1.0 (20) was used to run gene-wise one-tailed burden testing with a MAF<1% threshold. A 5% false discovery rate was used for multiple testing correction for 14 genes.

### Common variant association tests

SNPTEST 2.5.2 (46) was used to test for associated variants with MAF>1% across the region spanning the encoded gene. The first four covariates from the original GWAS were included (19). Bonferroni correction was used for 3,000 tests across the loci (*q*=1.66E-5).

### Plasmids

Myc-Flag-tagged *C1QTNF4* on the pCMV6 vector and the empty pCMV6 vector were used (OriGene). The mutant pCMV6-*C1QTNF4 C594G* (p.His198Gln) was generated by site-directed mutagenesis (Quikchange II XL; Stratagene) according the manufacturer instructions: mutagenic primer: 5’-GCGAGTGGTTGCTGCCGCGGCCC-3’ (Sigma Aldrich). The plasmids production was carried out in XL10-Gold Ultracompetent cells, isolated and purified using EndoFree Maxi Prep kit (Qiagen) and plasmid ORFs were confirmed by full Sanger sequencing (GATC-Biotech). The expression and secretion of the flagged proteins was confirmed by western blot on cell lysates and supernatants with monoclonal anti-FLAG antibody (clone M2; Sigma-Aldrich).

### Luciferase assays and TNF-induced programmed cell death

GloResponse NF-κB-RE-luc2P HEK293 cell line (Promega) and TNF-sensitive L929 fibrosarcoma cell line (ATCC) were cultured in Dulbecco’s Modified Eagle Medium (DMEM) supplemented with 10% fetal bovine serum (FBS) and 1% Penicillin/Streptomycin at 37°C, 5% CO_2_. HEK293 were seeded 24 hours before transfection in antibiotic free DMEM in 96 wells plate (2×10^4^ cells/well), transfected with either *C1QTNF4*, *C1QTNF4 C594G* or Empty Vector via Fugene HD (Promega). 48 hours after transfection the cell were left unstimulated or stimulated with TNFα 5 ng/ml (PeproTech) for 4 hours. Luciferase activity was assayed by One-Glo (Promega) on Berthold Orion luminometer, the values were normalized to cell viability measured by CellTiter Glo (Promega). L929 were challenged with TNFα 0.45 ng/ml and Actinomycin D 1μg/ml (R&D) for 24 hours in presence of C1QTNF4 or C1QTNF4 p.His198Gln containing media, cell viability was measured by CellTiter Glo.

### Size exclusion chromatography

Supernatants (750 μl) of HEK293 producing C1QTNF4 or C1QTNF4 p.His198Gln were buffer exchanged in PBS on Zeba Spin Desalting Columns (Thermo Fisher) and 0.5 mL loaded on an AKTA FPLC with a Superdex 200 10/300 GL column (GE Healthcare). Absorbance was normalized to the maximum peak of each sample.

### Data availability

WES data on 90 individuals – 30 parent-offspring trios – will be deposited at the European Genome-phenome Archive.

## Acknowledgements

The work leading to these results received funding from the European Union FP7 programme (grant agreement n^o^ 262055) via the European Sequencing and Genotyping Infrastructure (ESGI). Sequencing was performed by the SNP&SEQ Technology Platform in Uppsala, which is part of the National Genomics Infrastructure (NGI) hosted by Science for Life Laboratory in Sweden. This work was supported in part by the Swedish Research Council for Medicine and Health (grant n^o^ E0226301) and by the Knut and Alice Wallenberg Foundation (KAW 2011.0073). We thank Johanna Lagensjö and Olof Karlberg for assistance with sequencing. The research was funded/supported by the National Institute for Health Research (NIHR) Biomedical Research Centre based at Guy’s and St Thomas’ NHS Foundation Trust and King’s College London.

